# A searchable database of publications using fluorescent probes and flow cytometry to study antigen-specific B cells

**DOI:** 10.1101/2025.10.08.680531

**Authors:** Sunil Palani, Ally Xinyi Kong, Anna Buetow, Ashraf S. Yousif, Alex Say, Zhongchen Wang, Kuntal Biswas, Julia Blaszczyk, Monica J. Rodrigues-Jesus, Darlene R. Malavé Ramos, Vignesh Senthilkumar, Scott Chappell, Joshua F. E. Koenig, Justin J. Taylor

## Abstract

The study of antigen-specific B cells has resulted in important advances in all fields of immunology, the development of experimentally and/or clinically useful antibodies, and as a starting point for rationally designed vaccine antigens. A key innovation allowing for widespread study of antigen-specific B cells was the development of fluorescent antigen probes for use with flow cytometry. Initially these studies were mostly focused upon B cells specific for a variety of model antigens, but over the past decade focus has shifted towards the study of B cells specific for antigens from pathogens such as SARS-CoV-2, HIV, and Influenza virus. Importantly however, these types of approaches have been used for hundreds of different antigens and could be used for thousands more. Unfortunately, studies of B cells specific for an antigen of interest are not easily searchable on current publication databases since these assays are often a small portion of a larger publication. To overcome this, we built a searchable database of studies analyzing antigen-specific B cells by flow cytometry using fluorescent antigen probes that is located at www.immunology.virginia.edu/Taylor/Bcell/Database.php. Using this database, we assessed the number of publications per year revealing rapid growth in the use of this approach in recent years. While much of this rapid growth was focused upon the assessment of B cells specific for SARS-CoV-2, HIV-1, or Influenza virus, studies assessing B cells specific for hundreds of different antigens derived from numerous microbes, animals, plants, or other sources can be found in the database. Combined, the antigen-specific B cell database was built to facilitate identification of studies assessing these cells and for analysis of the field as a whole.

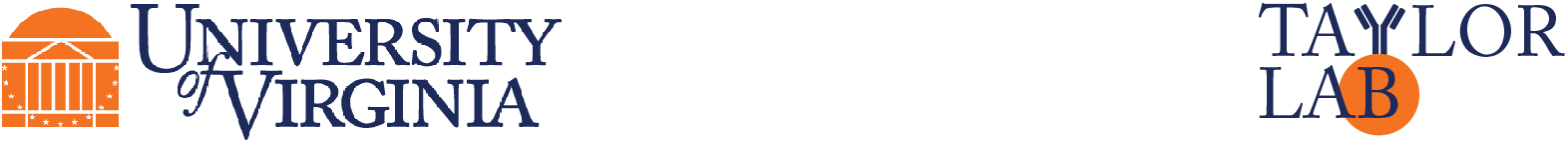

## DATABASE BUILDING

We begin this initiative in 2022 with the goal of building a reference list for an antigen tetramer validation publication (1). Shortly thereafter an early version of the list was shared on social media and got suggestions from other researchers for dozens of publications we had missed. From there we shifted to trying to identify all studies using fluorescent antigen probes analyze antigen-specific B cells by flow cytometry and began recording information about the types of probes used. This type of information is important because the type of probe used can dictate the sensitivity of the assay. For example, a monomeric probe may only identify B cells expressing higher affinity antibodies where a highly multimerized probe would be expected to allow assessment of cells unable to stably bind monomeric antigens (2). Likewise, probes using a bright fluorescent molecule with low background binding may better resolve a population of interest compared to a probe bound to dim fluorescent molecule, or one with high background binding to cells not specific for the antigen of interest. As such, we’ve included information on the type of probe as well as the fluorescent molecule used for each publication to aide users of the database.

Inclusion in the database was made solely upon the publication using fluorescent antigen probes to detect antigen-binding B cells by flow cytometry. Notably, we included studies using antigen probes that were covalently or non-covalently conjugated to fluorescent molecules, as well as studies in which cells were labeled with a non-fluorescent antigen followed by a fluorescent secondary reagent. We did not require that the publication validated that the probe-binding B cells were truly antigen-specific nor did we require display of the flow cytometry gating data in the article or supplement. While validation and data display are good practices, inclusion in the database allows users to determine the importance of those criteria rather than the database compilers. Likewise, database inclusion is not an endorsement of the quality of the findings or conclusions of the work. Researchers are best suited to make these types of qualitative decisions without biases added by the compilers. Additionally, the publications missing from the database are not intentional exclusions, just ones that the compilers have not yet discovered. We are hoping by making the database public users will contact us and share publications we have overlooked so they can be added.

## DATABASE INSIGHTS

As of September 2025, the database contains over 700 publications with the earliest being work coming from B cell researcher and flow cytometry pioneer Dr. Leonore Herzenberg’s group in 1972 (3). Adoption of these approaches was initially slow, with only 10 publications coming over the next 20 years (4-13) (**Fig. 1**). Publications of these types of studies increased from 1993-2010, with use rapidly afterwards, such that we have so far found over 80 publications in each of 2023 and 2024 (**Fig. 1**). While studies of antigen-specific B cells from humans, mice, and non-human primates are most common, B cells from rabbits (14-19), pigs (20-22), rats (23, 24), cows (25), and fish (26) have also been assessed, highlighting the flexibility of these approaches.

**Figure. 1.**
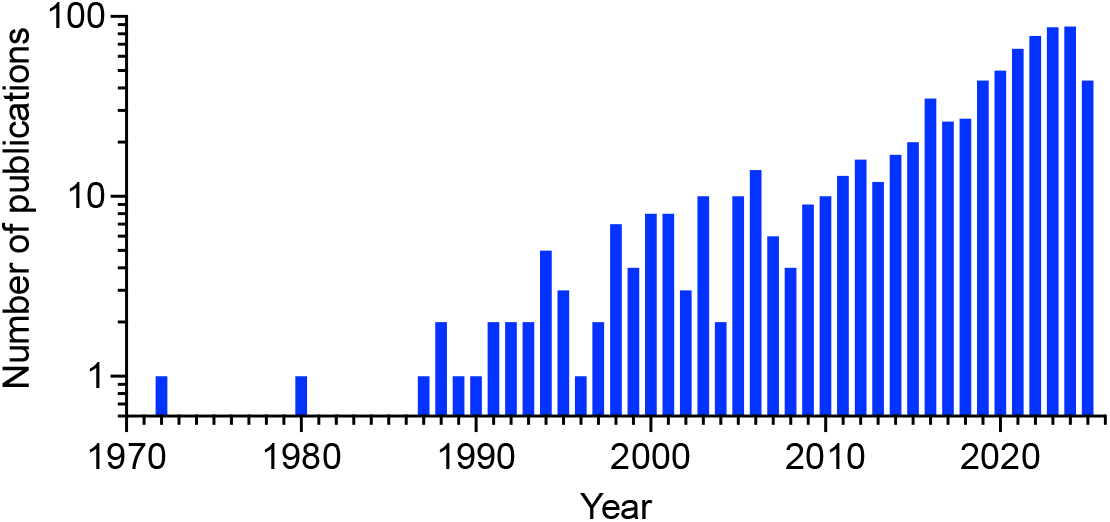
Number of studies per year using fluorescent antigen probes to assess antigen-specific B cells identified as of September 2025.

The papers in the database represent work from hundreds of labs and thousands of researchers. In fact, there are nearly 400 different last authors with multiple entries from most well-known B cell labs, but also many labs not known for B cell research. There is even more diversity in the nearly 600 different first authors, some of whom can also be found as last authors on later work from their independent labs.

One surprise upon building this database was the hundreds of different antigens studied derived from numerous microbes, animals, plants, or which were chemically synthesized (**Fig. 2**). In many cases we have found a single publication using a specific antigen, whereas there are other cases where unique variants of an antigen have been used in hundreds of studies.

**Figure. 2.**
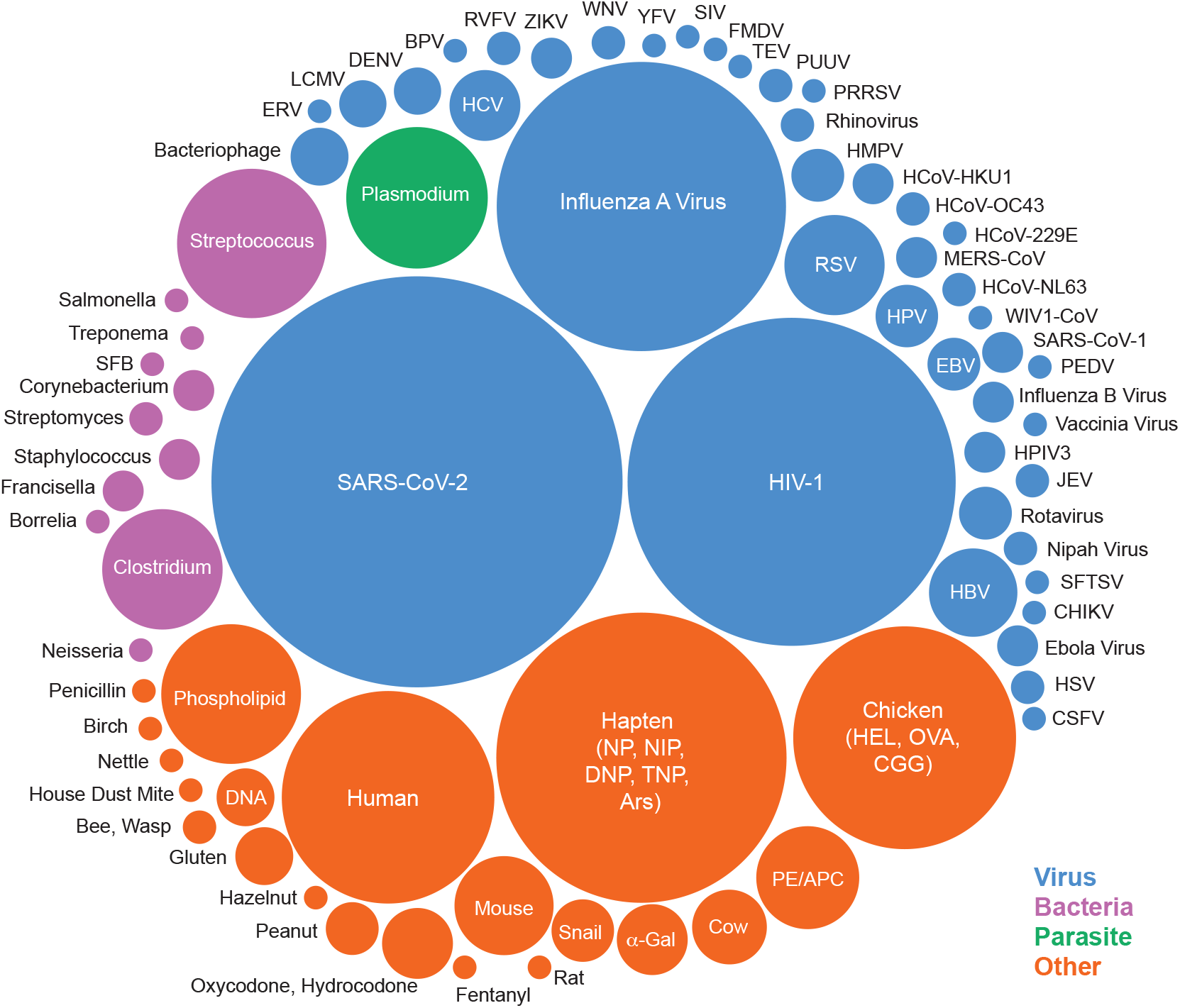
Overview of the usage different antigens used as probes in studies of antigen-specific B cells. This representation was generated based upon the usage of unique antigens used within each publications groups based upon the species the antigen was derived from, or type of antigen. The smallest circles represent one antigen used in a single publication whereas the largest circle represents 303 uses of several different antigens and antigen variants from SARS-CoV-2.

Also notable is the over 100 different journals in which antigen-specific B cell research has been published. Not surprisingly, the *Journal of Immunology* currently tops the list with over 80 publications, with *Journal of Experimental Medicine, Immunity, Cell Reports, Science Immunology, Frontiers in Immunology, Nature Immunology, Nature, Nature Communications, Cell, PNAS, Science, and The Journal of Clinical Investigation* also well represented on the list. Highlighting the breadth of research, there are many journals one might not expect studies of antigen-specific B cells to be found such as focused on microbiology, diabetes, parasitology, and pharmacology.

Below, we highlight a small fraction of the important work in the database.

### Studying model antigen-specific B cells

The use of model antigens has been critical in revealing fundamental properties of B cells. Amongst the model antigens used, the hapten 4-hydroxy-3-nitrophenyl acetyl (NP) and the related molecule 4-Hydroxy-3-iodo-5-nitrophenyl acetyl (NIP) have been most frequently utilized on the list since first used in a study by McHeyzer-Williams et. al. in 1991 (11). The list of first and last authors on papers studying NP/NIP-specific

B cell responses yields an impressive list of B cell researchers over the last four decades. One of the strengths of NP as model antigen is that it is not a protein but can be easily attached to a carrier protein like Keyhole Limpet Hemocyanin (KLH) as a source of peptides targeted by CD4^+^ helper T cells for the study of T-dependent responses or attached to a non-protein molecule like ficoll for the study of T-independent responses. In C57BL/6 mice, the response to NP is also dominated by B cells using VH186.2 and a lambda 1 light chain allowing for highly reproducible responses with well mapped out affinity maturation. The development of B cell receptor transgenic mice with high or low affinity for NP also helped define the role that naïve B cell affinity for antigen plays in B cell responses. Another advantage of NP as a model antigen is the commercial availability of fluorescent NP probes and NP-conjugated antigens for immunization.

Another frequently used model antigen is hen egg lysozyme (HEL). This was first used in a study by Goodnow et al. helped define mechanisms of B cell tolerance to self-antigen utilized a double-transgenic mouse model expressing HEL as a neo-self-antigen along with the high-affinity HEL-specific MD4 B cell receptor (BCR) (7). Shortly thereafter, studies by Hartley et al. analyzed the development of HEL-specific B cells upon HEL exposure in the bone marrow (27) and Cyster et al. explored peripheral tolerance development using the double-transgenic strategy and discovered that the exclusion of HEL-specific B cells from migrating into lymphoid follicles resulted in their premature death (28). Many of the same well-known researchers studying NP-specific B cells appear on the list of HEL-specific B cell studies. Importantly, not all studies of HEL-specific cells use B cells from the MD4 BCR transgenic mouse, which cannot undergo isotype class switching due to how this transgenic animal was generated. Instead, many studies analyzed B cells from the swHEL BCR transgenic mouse, which was generated in 2003 by Phan et. al. (29). The swHEL BCR transgenic expresses the same HEL-specific clonal sequence as the MD4 BCR transgenic but the swHEL antibody heavy chain sequence is knocked into the antibody heavy chain gene allowing for isotype class-switching (29, 30). The production of BCR knockin transgenics have become more accessible in recent years due to the emergence of CRISPR.

Fluorescent proteins commonly used for flow cytometry can themselves be used as model antigens. This approach was first used by Hayakawa et. al. in 1987 to study B cells specific for R-phycoerythrin (PE) (5), and used shortly thereafter in studies led by Schittek et. al. and Maruyama et. al. to define fundamental properties of memory B cells (9, 31). Fluorescent model antigens like PE and allophycocyanin (APC) are also helpful for identifying bonafide antigen-specific naïve B cells (32, 33) since the immunogens and detection reagents are the same whereas non-fluorescent antigens must be conjugated to a fluorescent molecule to be used as a detection reagent. Naïve B cells that bind such probes may be specific either for the target antigen or the fluorescent molecule. Approaches to exclude B cells of unwanted specificities is discussed in the *Types of probes* section.

### Studying microbe-specific B cells

In recent years the study of B cells specific for antigens derived from at least 60 microbes has eclipsed model antigens. Amongst these, the study of HIV-1-specific B cells has allowed for the identification of broadly neutralizing antibodies, tracking of vaccine responses, and study of fundamental B cell biology.

Studies by Moody et. al., Verkoczy et. al., Scheid et. al., and Wu et. al. in 2008 - 2010 (34-38) were the earliest assessments of HIV-1-specific B cells. Subsequently these types of approaches have been used extensively in human, mouse, and non-human primate studies by dozens of labs helping the field push towards a protective vaccine. In total, there are more than 150 studies looking at B cells specific for HIV-1 antigens. Many of these labs also were able to quickly pivot to the study of B cells specific for SARS-CoV-2, allowing for assessment of these responses remarkably early during the pandemic across the world. There are more than 300 publications assessing SARS-CoV-2-specific B cells by flow cytometry we have found so far, in part due to the commercial availability of antigens from this pathogen.

Identification of antigen-specific B cells is often hindered by binding of antigens to other molecules on B cells. This issue proved limiting for understanding humoral immune responses to influenza virus hemagglutinin (HA) and guiding vaccine development. The identification of authentic HA-specific B cells is complicated by the intrinsic sialic acid-binding capability of HA. A mutant variant of HA featuring a tyrosine to phenylalanine substitution at position 98 (Y98F) within the receptor binding site was initially employed by Whittle et. al. in 2014 (39) and widely adopted as an approach to overcome this issue. The Y98F mutation maintains the natural trimeric structure and conformational epitopes of HA while abolishing its capacity to bind sialic acid (40, 41) on non-influenza-specific B cells. By addressing the underlying issue of receptor-mediated non-specificity, this engineered probe has enabled high-fidelity detection and functional study of HA-specific B cells, boosting both basic immunology and translational vaccine research.

Plasmodium-specific B cells are another highly studied population with several dozen publications beginning with studies by Krishnamurty et. al. and Keitany et. al. in 2016 which used probes made of antigens from *Plasmodium chabaudi, Plasmodium falciparum*, and *Plasmodium yoelii* (42, 43). A 2021 study by Sutton et. al. of plasmodium-specific B cells focused on “atypical” memory B cells is particularly notable as the research into this phenotype continues to surge (44).

While most studies in the database focus on protein antigens, the ability to detect B cells specific for non-protein molecules like polysaccharides is essential for other types of infections and diseases. This is perhaps best highlighted by studies examining responses to pneumococcal polysaccharides after infection or vaccination. Beginning in 2012 studies by Khaskhely et. al., Leggat et. al., Iyer et. al., and Ohtola et. al. analyzed human B cells specific for pneumococcal polysaccharide using directly conjugated polysaccharide (45-51). More recently, a Tzovara et. al. study introduced a bead-based method in which biotinylated pneumococcal polysaccharide antigens from all 13 PCV13 vaccine serotypes were coupled to anti-biotin beads and subsequently labeled with PE-conjugated anti-biotin antibodies to generate fluorescent polysaccharide multimers (52, 53). A multimer-based approach was developed by Hoving et al. in 2023 where biotinylated polysaccharide antigens were conjugated to different combinations of fluorescent streptavidin, allowing combinatorial multimer staining in conjunction with a spectral flow cytometry panel to simultaneously characterize responses to 14 *Streptococcus pneumoniae* serotypes in vaccinated and unvaccinated donors (54).

### Studying self-antigen-specific B cells

Studying B cells specific for human or mouse antigens is important for understanding mechanisms underlying autoimmunity as well as transplant rejection. While B cells specific for numerous human and mouse antigens have been studied, a particularly interesting approach has used peptide:MHC monomers and tetramers typically generated for the study of antigen-specific T cells to assess MHC-specific B cells. A study by Mulder et. al. in 2003 was the first to assess human HLA-A2-specific B cells (55), and a study by Liu et. al. in 2005 was the first to assess H-2K^b^-specific B cells in mice (56). The availability of these types of tools from the NIH tetramer core at Emory and commercial sources can help facilitate these types of studies whereas researchers often need to produce and purify their own tools for other self-antigens.

### Studying allergen-specific B cells

To gain insight on these questions related to IgE-mediated allergies, researchers have begun to assess B cells specific for various allergens such as the peanut Ara h antigens. Work by Patil et. al. 2015 (57) and Hoh et. al. in 2016 (58), and more recently, 2023, studies Koenig et. al. (59) and Ota et. al. revealed important insights into the memory B cell subsets harboring IgE memory (60). Studies analyzing B cells specific for several other food and environmental allergens have been conducted and we expect many more in the coming years.

### Studying tumor antigen-specific B cells

Understanding the role of B cells in tumor control has been gaining prominence in recent years, and one long-standing question in the field is the specificity of B cells responding to tumors. These types of studies begin to appear in the last decade as researchers assess B cells specific for model tumor antigens like ovalbumin (OVA) and HEL by Guy et. al. and Wennhold et. al. (61, 62) and B cell responses to tumor antigens such as NY-ESO-1 by Wennhold et. al. and Schlosser et. al. (62, 63). A study by Wieland et. al. identified human papillomavirus (HPV)-specific B cell responses in HPV-positive head and neck squamous cell carcinoma (64). The authors observed the presence of HPV-specific B cells not only within the tumor microenvironment but also actively producing antibodies in a localized manner. The localization of tumor antigen-specific B cells holds significant potential in anti-tumor immunity through localized antigen presentation and/or antibody secretion, which is likely to be under rigorous assessment in the coming years.

### Studying antigen-specific B-1 cells

While most of the papers in the database focus on B-2 cell analysis, important studies of B-1 cells are also present. B-1 cells comprise a small but important subset of B cells predominantly found in the peritoneal and pleural cavities but present in low frequencies in other locations. B-1 cells also often display autoreactivity towards self-lipid antigens such as phosphatidylcholine (PtC), contributing to immune tolerance and modulating immune responses through cytokine secretion. Fluorescent PtC liposomes mimics PtC expression in cellular membranes and was first utilized in 1988 by Mercolino et. al. (6) and used in at least a dozen other studies since. This method enables phenotypic and functional analyses of B-1 cells and have shed light on their essential roles in immune homeostasis, tissue localization, natural IgM production, and defense against infection (65-68).

### Types of probes

There are numerous different types of probes used to detect antigen-specific B cells. Many studies use indirect staining with a non-fluorescent antigen followed by a secondary fluorescent reagent. Studies using antigens directly conjugated to a fluorochrome are also common, as are studies using antigens non-covalently multimerized or tetramerized using streptavidin. One difficulty in this type of analysis is that there are B cells specific for fluorochromes themselves, as mentioned above when discussing analyzing B cells specific for fluorochromes as convenient model antigens. Likewise, there are B cells specific for streptavidin or other molecules used to stabilize or multimerize antigens, as well as any tags added to aide in protein purification (69, 70). These unwanted B cells can be excluded by co-staining with antigens labeled with different fluorochromes as first shown by Townsend et. al. in 2001 (71), or by including a control “decoy” reagent to allow unwanted cells to be gated out developed by Taylor et. al. in 2012 (69, 70).

Other types of multimers have also be used for the analysis of antigen-specific B cells. An example of this is the use of virus-like particles (VLPs), which mimic the structure of native viruses and have been successfully employed in studies of rotavirus-specific B cells beginning in 2003 by Weitkamp et. al. (72) and bacteriophage QBeta-specific B cells beginning in 2011 by Hou et. al. (73). Similarly, Hepatitis B surface antigen multimers, which are produced naturally during infection, were first used in 2017 and 2018 studies led by Wennhold et. al. (62), Pinder et. al. (74), and Salimzadeh et. al. (75). Along the same lines, Scherer et. al. used fluorescent pseudoviruses to identify HPV-specific B cells beginning in 2014 (76).

With the development of high-throughput single cell sequencing technologies, the depth at which the field can interrogate B cells has quickly multiplied. Work in 2019 by Setliff et. al. first utilized antigens with a DNA barcode compatible with single cell sequencing applications (77), and made easier by using commercially available barcoded-streptavidin-PE/APC for tetramer production. Users of these barcoded tools may have trouble separating bonafide antigen-specific B cells from noise, a problem recent work by Wasdin et. al. helps to overcome (78). These types of approaches allow for high parameter analysis of antigen-specific B cells and will likely be used widely in the coming years.

## CONCLUSION

We hope this database helps researchers find relevant research and use probes for the study of antigen-specific B cells. Compiling this list has already proven valuable to the compilers, helping us to discover relevant work that aided technical and/or conceptual approaches. We hope others discover the same and encourage researchers to send us missing work so we can continue to make this a comprehensive resource.

## Abbreviations

a-Gal: Galactose-a-1,3-Galactose
APC: allophycocyanin
Ars: p-Azophenylarsonate
BCR: B cell receptor
BPV: Bovine papillomavirus
CGG: Chicken Gamma Globulin
CHIKV: Chikungunya virus
CSFV: Classical swine fever virus
DENV: Dengue virus
DNP: 2,4-Dinitrophenol
ERV: Endogenous Retrovirus
EBV: Epstein-Barr virus
FMDV: Foot-and-Mouth disease virus
HA: hemagglutinin
HBV: Hepatitis B Virus
HCV: Hepatitis C Virus
HCoV: Human coronavirus
HEL: Hen egg lysozyme
HIV: Human Immunodeficiency Virus
HMPV: Human Metapneumovirus
HPIV3: Human Parainfluenza Virus type 3
HPV: Human Papillomavirus
HSV: Herpes Simplex Virus
JEV: Japanese Encephalitis Virus
KLH: Keyhole Limpet Hemocyanin
LCMV: Lymphocytic choriomeningitis virus
MERS: Middle east respiratory syndrome
NIP: 4-Hydroxy-3-iodo-5-nitrophenyl acetyl
NP: 4-hydroxy-3-nitrophenyl acetyl
OVA: Ovalbumin
PE: R-phycoerythrin
PEDV: Porcine epidemic diarrhea virus
PRRSV: Porcine reproductive and respiratory syndrome virus
PtC: Phosphatidylcholine
PUUV: Puumala Hantavirus
RSV: Respiratory Syncytial Virus
RVFV: Rift Valley Fever Virus
SARS-CoV: Severe acute respiratory syndrome coronavirus
SFTSV: Severe Fever With Thrombocytopenia Syndrome Virus
SFB: Segmented Filamentous Bacteria
TBEV: Tick-borne Encephalitis Virus
TNP: 2,4,6-Trinitrophenyl
VLP: virus-like particle
YFV: Yellow Fever Virus
ZIKV: Zika Virus

## CONTRIBUTIONS

Justin J. Taylor conceived the database; added references and information about the antigens and probes; and wrote and edited this summary. Zhongchen (Seb) Wang, Vignesh Senthilkumar, Monica J. Rodrigues-Jesus, Ashraf S. Yousif, Alex Say, and Sunil Palani wrote and edited this this overview. Anna Buetow filled in missing information about the publications. Ally Kong added references and information about the antigens and probes. Scott Chappell developed the searchable database format and interface. Josh F. E. Koenig added references and information about the antigens/probes and wrote code to extract publication details.

## ACKNOWLEDGMENTS

We thank R. Guruge, H. Rodriguez, and K. Cummings for careful reading of the manuscript and suggestions regarding the figures; and D. Baumjohann, A. Broggi, K. Bruton, C. Castellanos, L. Chopp, C. Coelho, D. Glass, C. Hopp, S. Langel, N. Mohamed, K. O’Connor, G. Rizzuto, C. Sundling, L. Swadling, P. Wilson, F. Wimmers, R. Yefet, and T. Yewdell for suggesting missing publications in person, on social media, or email.

